# Leveraging multiple transcriptome assembly methods for improved gene structure annotation

**DOI:** 10.1101/216994

**Authors:** Luca Venturini, Shabhonam Caim, Gemy G Kaithakottil, Daniel L Mapleson, David Swarbreck

## Abstract

The performance of RNA-Seq aligners and assemblers varies greatly across different organisms and experiments, and often the optimal approach is not known beforehand. Here we show that the accuracy of transcript reconstruction can be boosted by combining multiple methods, and we present a novel algorithm to integrate multiple RNA-Seq assemblies into a coherent transcript annotation. Our algorithm can remove redundancies and select the best transcript models according to user-specified metrics, while solving common artefacts such as erroneous transcript chimerisms. We have implemented this method in an open-source Python3 and Cython program, Mikado, available at https://github.com/lucventurini/Mikado.

## Background

The annotation of eukaryotic genomes is typically a complex process which integrates multiple sources of extrinsic evidence to guide gene predictions. Improvements and cost reductions in the field of nucleic acid sequencing now make it feasible to generate a genome assembly and to obtain deep transcriptome data even for non-model organisms. However, for many of these species often there are only minimal EST and cDNA resources and limited availability of proteins from closely related species. In these cases, transcriptome data from high-throughput RNA sequencing (RNA-Seq) provides a vital source of evidence to aid gene structure annotation. A detailed map of the transcriptome can be built from a range of tissues, developmental stages and conditions, aiding the annotation of transcription start sites, exons, alternative splice variants and polyadenylation sites.

Currently, one of the most commonly used technology for RNA-Seq is Illumina sequencing, which is characterised by extremely high throughput and relatively short read lengths. Since its introduction, numerous algorithms have been proposed to analyse its output. Many of these tools focus on the problem of assigning reads to known genes to infer their abundance [1–4], or of aligning them to their genomic locus of origin [5–7]. Another challenging task is the reconstruction of the original sequence and genomic structure of transcripts directly from sequencing data. Many approaches developed for this purpose leverage genomic alignments [8–11], although there are alternatives based instead on *de novo* assembly [9, 12, 13]. While these methods focus on how to analyse a single dataset, related research has examined how to integrate assemblies from multiple samples. While some researchers advocate for merging together reads from multiple samples and assembling them jointly [9], others have developed methods to integrate multiple assemblies into a single coherent annotation [8, 14].

The availability of multiple methods has generated interest in understanding the relative merits of each approach [15–17]. The correct reconstruction of transcripts is often hampered by the presence of multiple isoforms at each locus and the extreme variability of expression levels, and therefore in sequencing depth, within and across gene loci. This variability also affects the correct identification of transcription start and end sites, as sequencing depth typical drops near the terminal ends of transcripts. The issue is particularly severe in compact genomes, where genes are clustered within small intergenic distances. Further, the presence of tandemly duplicated genes can lead to alignment artefacts that then result in multiple genes being incorrectly reconstructed as a fused transcript. As observed in a comparison performed by the RGASP consortium [18], the accuracy of each tool depends on how it corrects for each of these potential sources of errors. However, it also depends on other external factors such as the quality of the input sequencing data as well as on species-dependent characteristics, such as intron sizes and the extent of alternative splicing. It has also been observed that no single method consistently delivers the most accurate transcript set when tested across different species. Therefore, none of them can be determined *a priori* as the most appropriate for a given experiment [19]. These considerations are an important concern in the design of genome annotation pipelines, as transcript assemblies are a common component of evidence guided approaches that integrate data from multiple sources (e.g. cDNAs, protein or whole genome alignments). The quality and completeness of the assembled transcript set can therefore substantially impact on downstream annotation.

Following these studies, various approaches have been proposed to determine the best assembly using multiple measures of assembly quality [19, 20] or to integrate RNA-Seq assemblies generated by competing methods [21–23]. In this study we show that alternative methods not only have different strengths and weaknesses, but that they also often complement each other by correctly reconstructing different subsets of transcripts. Therefore, methods that are not the best overall might nonetheless be capable of outperforming the “best” method for a sub-set of loci. An annotation project typically integrates datasets from a range of tissues or conditions, or may utilise public data generated with different technologies (e.g. Illumina, PacBio) or sequencing characteristics (e.g. read length, strand specificity, ribo-depletion); in such cases, it is not uncommon to produce at least one set of transcript assemblies for each of the different sources of data, assemblies which then need to be reconciled. To address these challenges, we developed MIKADO, an approach to integrate transcript assemblies. The tool defines loci, scores transcripts, determines a representative transcript for each locus, and finally returns a set of gene models filtered to individual requirements, for example removing transcripts that are chimeric, fragmented or with short or disrupted coding sequences. Our approach was shown to outperform both standalone methods and those that combine assemblies, by returning more transcripts reconstructed correctly and less chimeric and unannotated genes.

## Results and discussion

Assessment of RNA-Seq based transcript reconstruction methods

We evaluated the performance of four commonly utilised transcript assemblers: Cufflinks, StringTie, CLASS2 and Trinity. Their behaviour was assessed in four species, using as input data RNA-Seq reads aligned with two alternative leading aligners, TopHat2 and STAR. In total, we generated 32 different transcript assemblies, eight per species. In line with the previous RGASP evaluation, we performed our tests on the three metazoan species of *Caenhorabditis elegans, Drosophila melanogaster* and *Homo sapiens*, using RNA-Seq data from that study as input. We also added to the panel a plant species, *Arabidopsis thaliana*, to assess the performance of these tools on a non-metazoan genome. Each of these species has undergone extensive manual curation to refine gene structures, and moreover, these annotations exhibit very different gene characteristics in terms of their proportion of single exon genes, average intron lengths and number of annotated transcripts per gene (Supplementary Table ST1). Similar to previous studies [18, 24], we based our initial assessment on real rather than simulated data, to ensure we captured the true characteristics of RNA-Seq data. Prediction performance was benchmarked against the subset of annotated transcripts with all exons and introns (minimum 1X coverage) identified by at least one of the two RNA-Seq aligners.

The number of transcripts assembled varied substantially across methods, with StringTie and Trinity generally reconstructing a greater number of transcripts (Supplementary Figure SF1). Assembly with Trinity was performed using the genome guided denovo method, where RNA-Seq reads are first partitioned into loci ahead of de-novo assembly. This approach is in contrast to the genome guided approaches employed by the other assemblers that allow small drops in read coverage to be bridged and enable the exclusion of retained introns and other lowly expressed fragments. As expected Trinity annotated more fragmented loci, with a higher proportion of mono-exonic genes (Supplementary Figure SF1).

Accuracy of transcript reconstruction was measured using recall and precision. For any given feature (nucleotide, exon, transcript, gene), recall is defined as the percentage of correctly predicted features out of all expressed reference features, whereas precision is defined as the percentage of all features that correctly match a feature present in the reference. In line with previous evaluations, we found that accuracy varied considerably among methods, with clear trade-offs between recall and precision (Supplementary Figure SF2). For instance, CLASS2 emerged as the most precise of all methods tested, but its precision came at the cost of reconstructing less transcripts overall. In contrast, Trinity and StringTie often outperformed the recall of CLASS2, but were also much more prone to yield erroneous transcripts (Supplementary Figure SF2, SF3). Notably, the performance and the relative ranking of the methods differed among the four species (Table 1). We found CLASS2 and StringTie to be overall the most accurate (with either aligner), however exceptions were evident. For instance, the most accurate method in D. melanogaster (CLASS2 in conjunction with Tophat alignments) performed worse than any other tested method in *A. thaliana*. The choice of RNA-Seq aligner also substantially impacted assembly accuracy, with clear differences between the two when used in conjunction with the same assembler.

**Table 1:** Cumulative z-score for each method aggregating individual z-scores based on base, exon, intron, intron chain, transcript and gene F1 score (top ranked method in light gray, bottom ranked method in dark gray).

| Method | *A. thaliana* |  | *C. elegans* |  | *D. melanogaster* |  | *H. sapiens* |  | All methods |  |
| --- | --- | --- | --- | --- | --- | --- | --- | --- | --- | --- |
|  | Z-score | Rank | Z-score | Rank | Z-score | Rank | Z-score | Rank | Z-score | Rank |
| CLASS2 (STAR) | 7.627 | 1 | 7.309 | 1 | -3.310 | 6 | 5.258 | 1 | 16.884 | 1 |
| StringTie (TopHat2) | 0.584 | 4 | 5.502 | 3 | 6.612 | 2 | 3.199 | 3 | 15.897 | 2 |
| CLASS2 (TopHat2) | -5.542 | 8 | 6.698 | 2 | 9.314 | 1 | 4.998 | 2 | 15.738 | 3 |
| StringTie (STAR) | 2.621 | 3 | -2.197 | 4 | 1.587 | 3 | 2.991 | 4 | 5.001 | 4 |
| Cufflinks (STAR) | 2.716 | 2 | -2.306 | 5 | -1.730 | 5 | 1.037 | 5 | -0.283 | 5 |
| Cufflinks (TopHat2) | -0.526 | 5 | -5.363 | 8 | -1.504 | 4 | -0.993 | 6 | -8.386 | 6 |
| Trinity (STAR) | -4.120 | 7 | -5.079 | 7 | -4.762 | 7 | -3.417 | 7 | -17.458 | 7 |
| Trinity (TopHat2) | -3.280 | 6 | -4.833 | 6 | -6.206 | 8 | -13.073 | 8 | -27.392 | 8 |

Across the four species and depending on the aligner used, 22 to 35% of transcripts could be reconstructed by any combination of aligner and assembler (Supplementary Table ST2). However, some genes were recovered only by a subset of the methods (Supplementary Table ST2), with on average 5% of the genes being fully reconstructable only by one of the available combinations of aligner and assembler.

Taking the union of genes fully reconstructed by any of the methods shows that an additional 14.92-19.08% of genes could be recovered by an approach that would integrate the most sensitive assembly with less comprehensive methods. This complementarity manifests as well in relation to genes missed by any particular method: while each approach failed to reconstruct several hundred genes on average, the majority of these models could be fully or partially reconstructed by an alternative method (Supplementary Figure SF3a). Another class of error are artifactual fusion/chimeric transcripts that chain together multiple genes. These artefacts usually arise from an incorrect identification of start and end sites during transcript reconstruction - an issue which appears most prominently in compact genomes with smaller intergenic distances [9]. Among the methods tested, Cufflinks was particularly prone to this class of error, while Trinity and CLASS2 assembled far fewer such transcripts. Again, alternative methods complemented each other, with many genes fused by one assembler being reconstructed correctly by another approach (Supplementary Figure SF3b). Finally, the efficiency of transcript reconstruction depends on coverage, a reflection of sequencing depth and expression level. Methods in general agree on the reconstruction of well-expressed genes, while they show greater variability with transcripts that are present at lower expression levels. Even at high expression levels, though, only a minority of genes can be reconstructed correctly by every tested combination of aligner and assembler (Supplementary Figure SF4). Our results underscore the difficulty of transcript assembly and highlight advantageous features of specific methods. A naive combination of the output of all methods would yield the greatest sensitivity, but at the cost of a decrease in precision as noise from erroneous reconstructions accumulates. Indeed, this is what we observe: in all species, while the recall of the naive combination markedly improves even upon the most sensitive method, the precision decreases (Supplementary Figure SF2). As transcript reconstruction methods exhibit idiosyncratic strengths and weaknesses an approach that can integrate multiple assemblies can potentially lead to a more accurate and comprehensive set of gene models.

### Overview of the Mikado method

Mikado provides a framework for integrating transcripts from multiple sources into a consolidated set of gene annotations. Our approach assesses, scores (based on user configurable criteria) and selects transcripts from a larger transcript pool, leveraging transcript assemblies generated by alternative methods or from multiple samples and sequencing technologies. The software takes as input transcript structures in standard formats such as GTF and GFF3, with optionally BLAST similarity scores or a set of high quality splice junctions, and produces a polished annotation and a rich set of metrics for each transcript. The software is written in python3 and Cython, and extensive documentation is available from https://github.com/lucventurini/mikado.

Mikado is composed of three core programs (prepare, serialise, pick) executed in series. The Mikado prepare step validates and standardizes transcripts, removing exact duplicates and artefactual assemblies such as those with ambiguous strand orientation (as indicated by canonical splicing). During the Mikado serialise step, data from multiple sources are brought together inside a common database. Mikado by default analyses and integrates three types of data: open-reading frames (ORFs) currently identified via TransDecoder, protein similarity derived through BLASTX or Diamond and high quality splice junctions identified using tools such as Portcullis [25] or Stampy [26]. The selection phase (Mikado pick) groups transcripts into loci and calculates for each transcript over fifty numerical and categorical metrics based on either external data (e.g. BLAST support) or intrinsic qualities relating to CDS, exon, intron or UTR features (summarised in Supplementary Table ST3). While some metrics are inherent to each transcript (e.g. the cDNA length), others depend on the context of the locus the transcript is placed in. A typical example would be the proportion of introns of the transcript relative to the number of introns associated to the genomic locus. Such values are dependent on the loci grouping, and can change throughout the computation, as transcripts are moved into a different locus or filtered out. Notably, the presence of open reading frames is used in conjunction with protein similarity to identify and resolve fusion transcripts. Transcripts with multiple ORFs are marked as candidate false-fusions; homology to reference proteins is then optionally used to determine whether the ORFs derive from more than one gene. If the fusion event is confirmed, the transcript is split into multiple transcripts (Figure 1).

**Figure 1:**
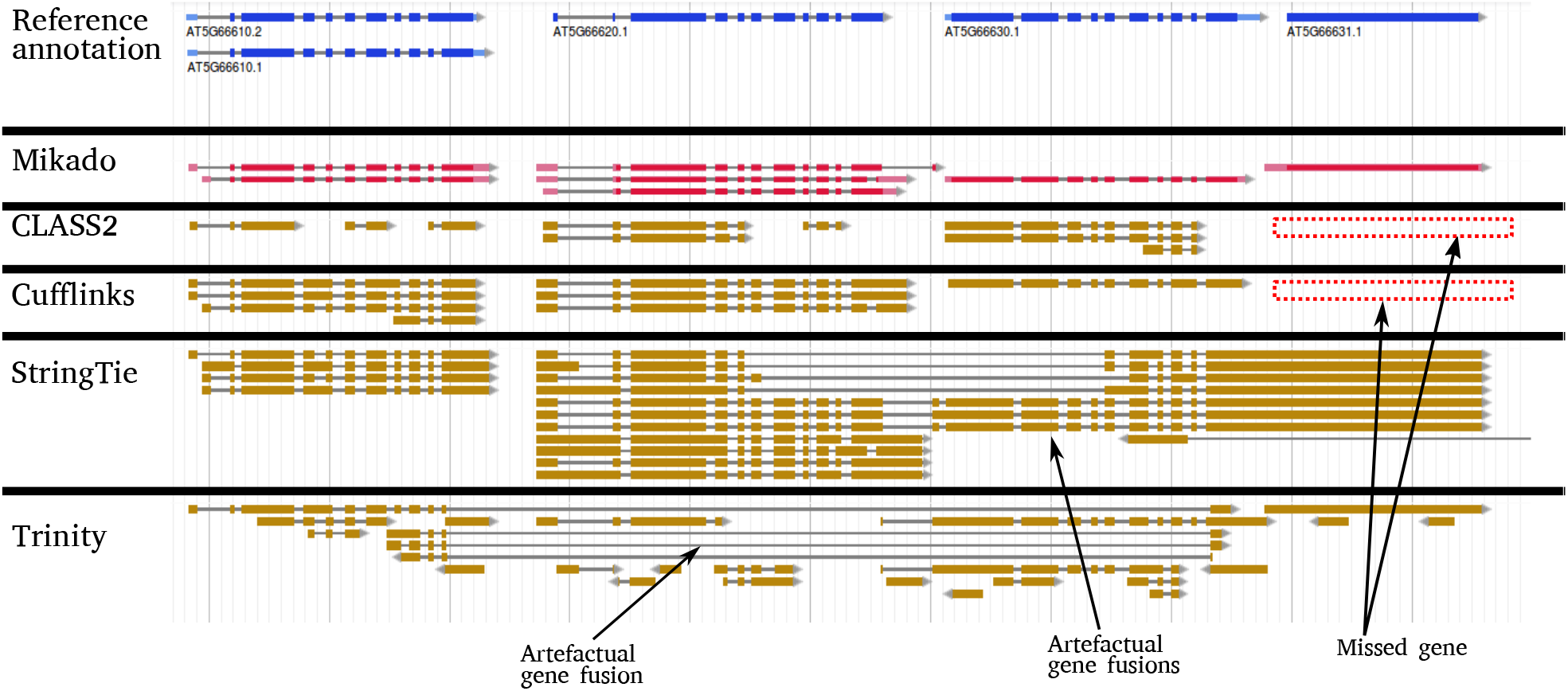
Example region. The algorithm employed by Mikado is capable of solving complex loci, with multiple potential assemblies. This locus in *A. thaliana* is particularly challenging as an ancestral gene in the locus tandemly duplicated into the current AT5G66610, AT5G66620 and AT5G66630 genes. Due to these difficulties, no single assembler was capable of reconstructing correctly all loci. For instance, Trinity was the only method which correctly assembled AT5G66631, but it failed to reconstruct correctly any other transcript. The reverse was true for Cufflinks, which correctly assembled the three duplicated genes, but completely missed the monoexonic AT566631. By choosing between different alternative assemblies, Mikado was capable to provide an evidence-based annotation congruent to the TAIR10 models.

To determine the primary transcript at a locus, Mikado assigns a score for each metric of each transcript, by assessing its value relatively to all other transcripts associated to the locus. Once the highest scoring transcript for the group has been selected, Mikado will exclude all transcripts which are directly intersecting it, and if any remain, iteratively select the next best scoring transcripts pruning the graph until all nonintersecting transcripts have been selected. This iterative strategy ensures that no locus is excluded if e.g. there are unresolved read-through events that would connect two or more gene loci. Grouping and filtering happen in multiple sequential phases, each defined by different rules for clustering transcripts into loci (see methods). The process is controlled by a configuration file that determines desirable gene features, allowing the user to define criteria for transcript filtering and scoring as well as specifying minimum requirements for potential alternative splicing events.

We also developed a Snakemake-based pipeline, Daijin, in order to drive Mikado, including the calls to external programs to calculate ORFs and protein homology. Daijin works in two independent stages, assemble and mikado. The former stage enables transcript assemblies to be generated from the read datasets using a choice of read alignment and assembly methods. In parallel, this part of the pipeline will also calculate reliable junctions for each alignment using Portcullis. The latter stage of the pipeline drives the steps necessary to execute Mikado, both in terms of the required steps for our program (prepare, serialise, pick) and of the external programs needed to obtain additional data for the picking stage (currently, homology search and ORF detection). A summary of the Daijin pipeline is reported in Figure.

### Performance of Mikado

To provide a more complete assessment we evaluated the performance of Mikado on both simulated and real data. While real data represents more fully the true complexity of the transcriptome simulated data generates a known set of transcripts to enable a precise assessment of prediction quality. For our purposes, we used SPANKI to simulate RNA-Seq reads for all four species, closely matching the quality and expression profiles of the corresponding real data. Simulated reads were aligned and assembled following the same protocol that was used for real data, above. Mikado was then used to integrate the four different transcript assemblies for each alignment.

Across the four species and on both simulated and real data, Mikado was able to successfully combine the different assemblies, obtaining a higher accuracy than most individual tools in isolation. Compared with the best overall combination, CLASS2 on STAR alignments, Mikado improved the accuracy on average by 6.58% and 9.23% on simulated and real data at the transcript level, respectively (Figure 3 and Additional File 2). Most of this improvement accrues due to an improved recall rate without significant losses on precision. We register a single exception, on *H. sapiens* simulated data, due to an excess of intronic gene models which pervade the assemblies of all other tools. On simulated data, CLASS2 is able to detect these models and exclude them, most probably using its refined filter on low-coverage regions [11]; however, this increase in precision is absent when using TopHat2 as an aligner and on real data. Aside from the accuracy in correctly reconstructing transcript structures, in our experiments, merging and filtering the assemblies proved an effective strategy to produce a comprehensive transcript catalogue: Mikado consistently retrieved more loci than the most accurate tools, while avoiding the sharp drop in precision of more sensitive methods such as e.g. Trinity (Figure 3b). Finally, Mikado was capable to accurately identify and solve cases of artefactual gene fusions, which mar the performance of many assemblers. As this kind of error is more prevalent in our real data, the increase in precision obtained by using Mikado was greater using real rather than simulated data.

**Figure 2:**
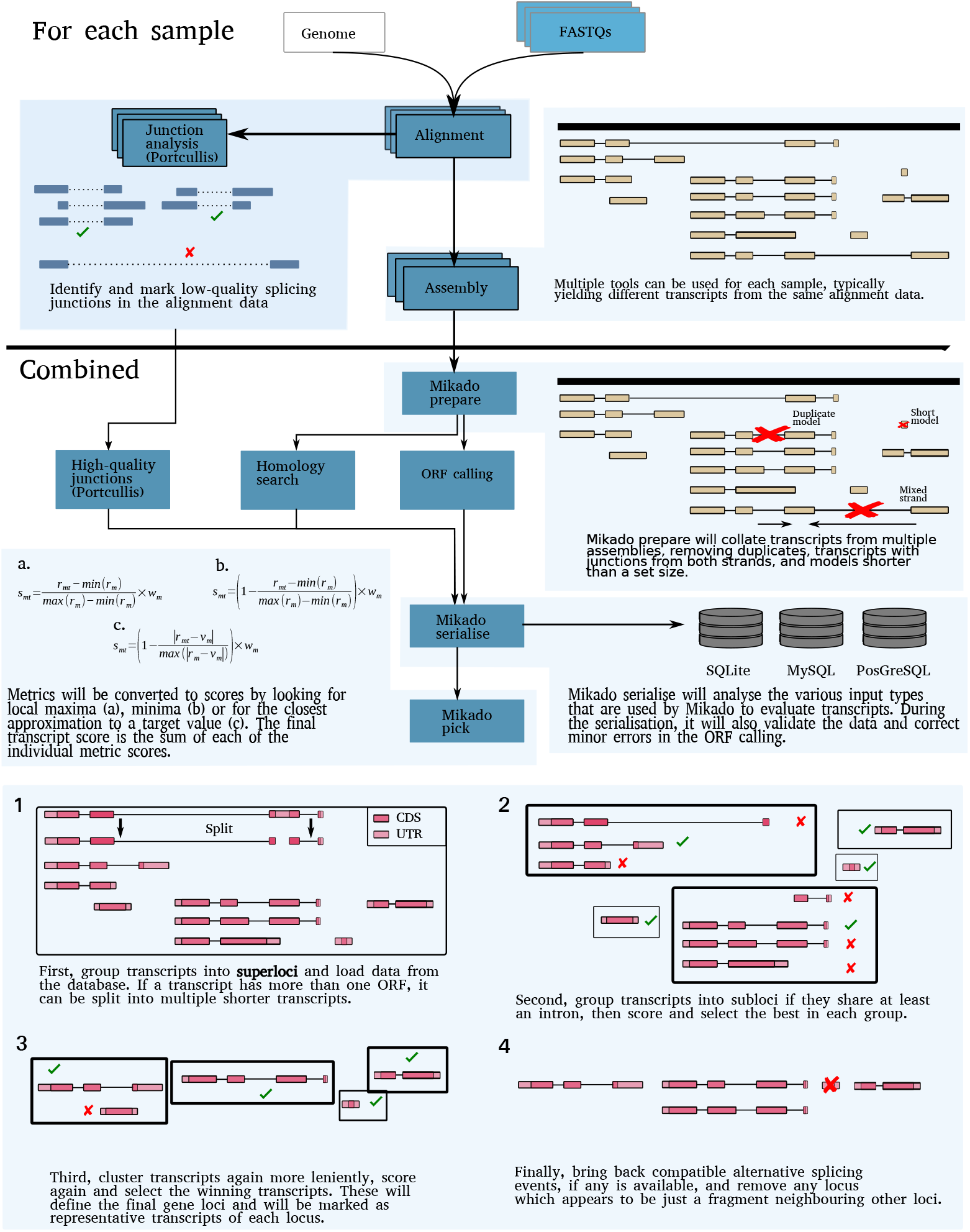
Schematic representation of the Mikado workflow.

**Figure 3:**
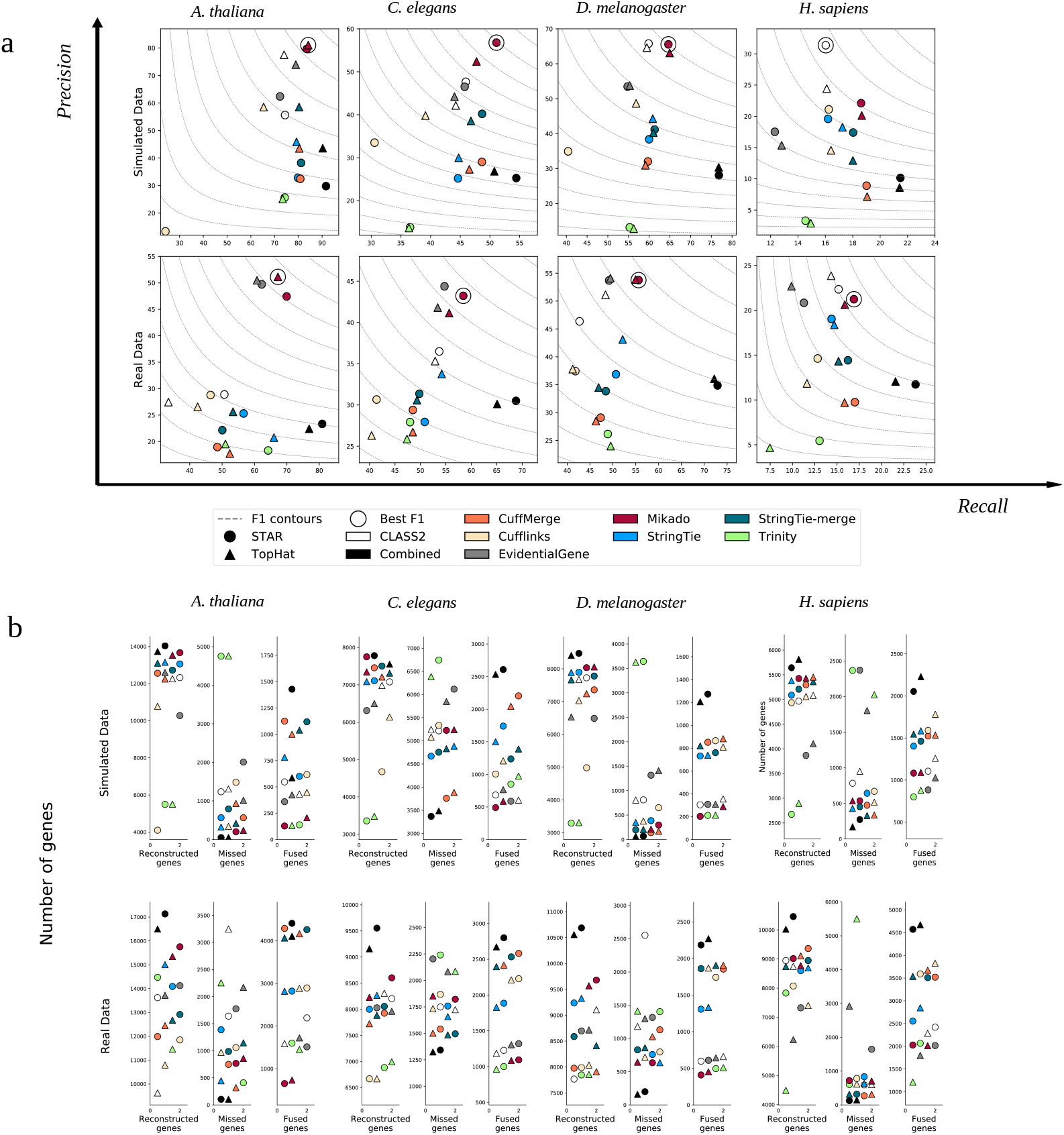
Performance of Mikado on simulated and real data. **a** We evaluated the performance of Mikado using both simulated data and the original real data. The method with the best transcript-level F1 is marked by a circle. **b** Number of reconstructed, missed and chimeric genes in each of the assemblies. Notice the lower level of chimeric events in simulated data.

We further assessed the performance of Mikado in comparison with three other methods that are capable of integrating transcripts from multiple sources: CuffMerge [27], StringTie-merge [14] and Evidential-Gene [23, 28]. CuffMerge and StringTie-merge perform a meta-assembly of transcript structures, without considering ORFs or homology. In contrast, EvidentialGene is similar to Mikado in that it classifies and selects transcripts, calculating ORFs and associated quality metrics from each transcript to inform its choice. In our tests, Mikado consistently performed better than alternative combiners, in particular when compared to the two meta-assemblers. The performance of StringTie-merge and CuffMerge on simulated data underscored the advantage of integrating assemblies from multiple sources as both methods generally improved recall over input methods. However, this was accompanied by a drop in precision, most noticeably for CuffMerge, as assembly artefacts present in the input assemblies accumulated in the merged dataset. In contrast, the classification and filtering based approach of EvidentialGene led to a more precise dataset, but at the cost of a decrease in recall. Mikado managed to balance both aspects, thus showing a better accuracy than any of the alternative approaches (*A. thaliana* +6.24%, *C. elegans* +7.66%, *D. melanogaster* +9.48%, *H. sapiens* +4.92% F1 improvement over the best alternative method). On real and simulated data, Mikado and EvidentialGene generally performed better than the two meta-assemblers, with an accuracy differential that ranged from moderate in *H. sapiens* (1.67 to 4.32%) to very marked in *A. thaliana* (14.87 to 29.58%). An important factor affecting the accuracy of the meta-assemblers with real data is the prevalence of erroneous transcript fusions that can result from incorrect read alignment, genomic DNA contamination or bona fide overlap between transcriptional units. Both StringTie-merge and CuffMerge were extremely prone to this type of error, as across the four species they generated on average 2.39 times the number of fusion genes compared to alternative methods (Figure 3b). Between the two selection based methods, Evidential-Gene performed similarly to Mikado on real data but much worse on simulated data: its accuracy was on average 2 points lower than Mikado on real data, and 8.13 points lower in the simulations. This is mostly due to a much higher precision differential between the two methods in simulated data, with Mikado much better than EvidentialGene on this front (+8.95% precision on simulated data).

### Filtering lenient assemblies

Although our tests have been conducted using default parameters for the various assemblers, these parameters can be adjusted to alter the balance between precision and sensitivity according to the goal of the experiment. In particular, three of the assemblers we tested provide a parameter to filter out alternative isoforms with a low abundance. This parameter is commonly referred to as “minimum isoform fraction”, or MIF, and sets for each gene a minimum isoform expression threshold relative to the most expressed isoform. Only transcripts whose abundance ratio is greater than the MIF threshold are reported. Therefore, lowering this parameter will yield a higher number of isoforms per locus, retaining transcripts that are expressed at low levels and potentially increasing the number of correctly reconstructed transcripts. This improved recall is obtained at the cost of a drop in precision, as more and more incorrect splicing events are reported (Supplementary Figure 4). Mikado can be applied on top of these very permissive assemblies to filter out spurious splicing events. In general, filtering with Mikado yielded transcript datasets that are more precise than those produced by the assemblers at any level of chosen MIF, or even when comparing the most relaxed MIF in Mikado with the most conservative in the raw assembler output (Figure 4).

**Figure 4:**
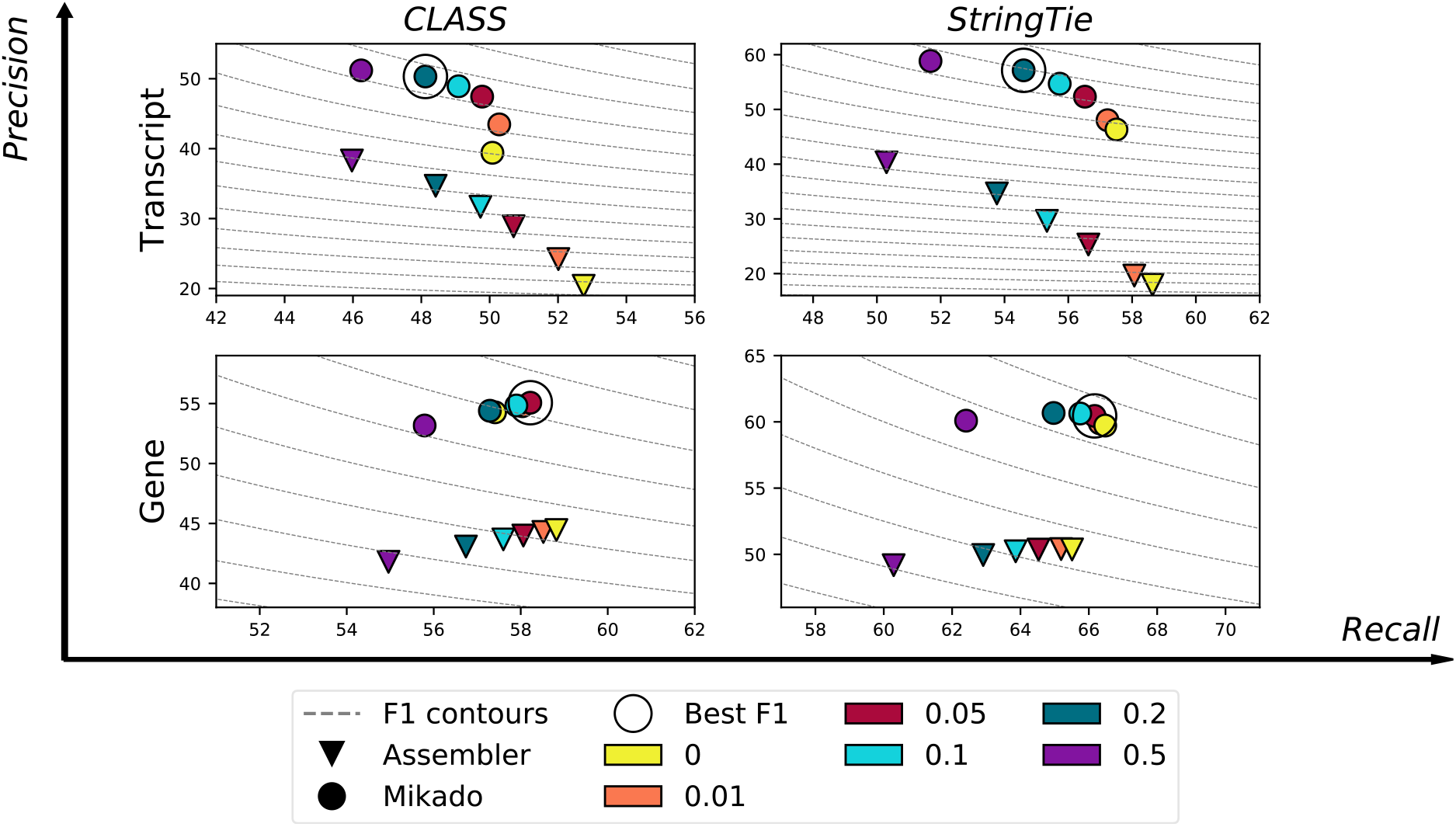
Performance of Mikado while varying the Minimum Isoform Fraction parameter. Precision/recall plot at the gene and transcript level for CLASS and StringTie at varying minimum isoform fraction thresholds in *A. thaliana*, with and without applying Mikado. Dashed lines mark the F1 levels at different precision and recall values. CLASS sets MIF to 5% by default (red), while StringTie uses a slightly more stringent default value of 10% (cyan).

### Multi-sample transcript reconstruction

Unravelling the complexity of the transcriptome requires assessing transcriptional dynamics across many samples. Projects aimed at transcript discovery and genome annotation typically utilize datasets generated across multiple tissues and experimental conditions to provide a more complete representation of the transcriptional landscape. Even if a single assembly method is chosen, there is often a need to integrate transcript assemblies constructed from multiple samples. StringTie-merge, CuffMerge and the recently published TACO [29] have been developed with this specific purpose in mind. The meta-assembly approach of these tools can reconstruct full-length transcripts when they are fragmented in individual assemblies, but as observed earlier, it is prone to creating fusion transcripts. TACO directly addresses this issue with a dedicated algorithmic improvement, ie change-point detection. This solution is based on fusion transcripts showing a dip in read coverage in regions of incorrect assembly; this change in coverage can then be used to identify the correct breakpoint. A limitation of the implementation in TACO is that it requires expression estimates to be encoded in the input GTFs, and some tools do not provide this information.

To assess the performance of Mikado for multi sample reconstruction, we individually aligned and assembled the twelve *A. thaliana* seed development samples from PRJEB7093, using the four single-sample assemblers described previously. The collection of twelve assemblies per tool was then integrated into a single set of assemblies, using different combiners. StringTie-merge and TACO could not be applied to the Trinity dataset, as they both require embedded expression data in the GTF files, which is not provided in the Trinity output. In line with the results published in the TACO paper [29], we observed a high rate of fusion events in both StringTie-merge and CuffMerge results (Figure 5b), which TACO reduced. However, none of these tools performed as well as EvidentialGene or Mikado, either in terms of accuracy, or in avoiding gene fusions (Figure 5). Mikado achieved the highest accuracy irrespective of the single sample assembler used, with an improvement in F1 over the best alternative method of +8.25% for Cufflinks assemblies, +2.23% in StringTie, +0.95% with CLASS2 and +6.65% for Trinity.

**Figure 5:**
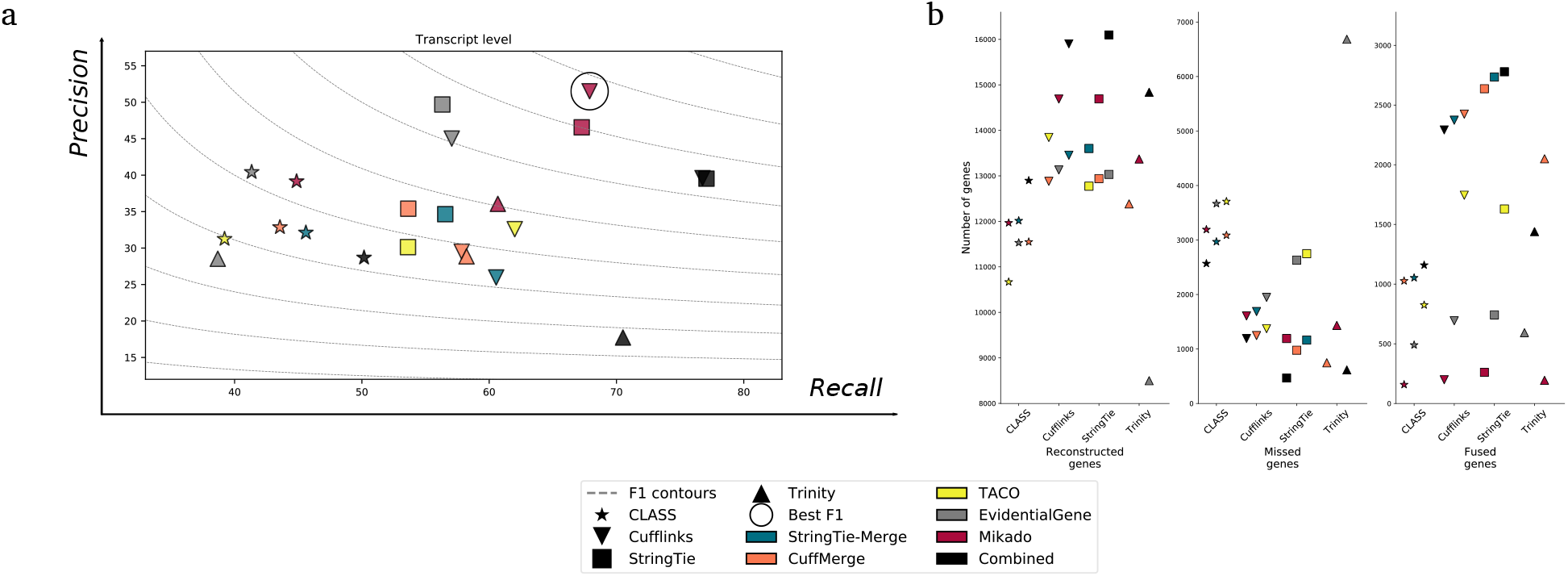
Integrating assemblies coming from multiple samples. **a** Mikado performs consistently better than other merging tools. StringTie-merge and TACO are not compatible with Trinity results and as such have not been included in the comparison. **b** Rate of recovered, missed, and fused genes for all the assembler and combiner combinations.

Transcript assemblies are commonly incorporated into evidence-based gene finding pipelines, often in conjunction with other external evidence such as cross species protein sequences, proteomics data or synteny. The quality of transcript assembly can therefore potentially impact on downstream gene prediction. To test the magnitude of this effect, we used the data from these experiments on *A. thaliana* to perform gene prediction with the popular MAKER annotation pipeline, using Augustus with default parameters for the species as a gene predictor. Our results (Supplementary Figure SF5) show that, as expected, an increased accuracy in the transcriptomic dataset leads to an increased accuracy in the final annotation. Importantly, MAKER was not capable of reducing the prevalence of gene fusion events present in the transcript assemblies. This suggests that *ab initio* Augustus predictions utilized by MAKER do not compensate for incorrect fusion transcripts that are provided as evidence, and stress the importance of pruning these mistakes from transcript assemblies before performing an evidence-guided gene prediction.

### Expansion to long read technologies

Short read technologies, due to their low per-base cost and extensive breadth and depth of coverage, are commonly utilised in genome annotation pipelines. However, like the previous generation Sanger ESTs, their short size requires the use of sophisticated methods to reconstruct the structure of the original RNA molecules. Third-generation sequencing technologies promise to remove this limitation, by generating full-length cDNA sequences. These new technologies currently offer lower throughput and are less cost effective, but have in recent studies been employed alongside short read technologies to define the transcriptome of species with large gene content [30, 31].

We tested the complementarity of the two technologies by sequencing two samples of a standard human reference RNA library with the leading technologies for both approaches, Illumina HiSeq for short-reads (250 bp, paired-end reads) and the Pacific Bioscience IsoSeq protocol for long reads. Given the currently higher per-base costs of long-read sequencing technologies, read coverage is usually much lower than for short read sequencing. We found many genes to be reconstructed by both platforms, but as expected given the lower sequencing depth there was a clear advantage for the Illumina dataset on genes with expression lower than 10 TPM (Supplementary Figures SF6a and SF6b). We verified the feasibility of integrating the results given by the different sequencing technologies by combining the long reads with the short read assemblies, either simply concatenating them, or by filtering them with EvidentialGene and Mikado (Supplementary Figure SF7). An advantage of Mikado over the two alternative approaches is that it allows to prioritise PacBio reads over Illumina assemblies, by giving them a slightly higher base score. In this analysis, we saw that even PacBio data on its own might require some filtering, as the original sample contains a mixture of whole and fragmented molecules, together with immature transcripts. Both Mikado and EvidentialGene are capable of identifying mature coding transcripts in the data, but Mikado shows a better recall and general accuracy rate, albeit at the cost of some precision. However, Mikado performed much better than Evidential-Gene in filtering either the Illumina data on its own, or the combination of the two technologies. Although the filtering inevitably loses some of the real transcripts, the loss is compensated by an increased overall accuracy. Mikado performed better in this respect than EvidentialGene, as the latter did not noticeably improve in accuracy when given a combination of PacBio and Illumina data, rather than the Illumina data alone.

## Conclusions

Transcriptome assembly is a crucial component of genome annotation workflows, however, correctly reconstructing transcripts from short RNA-Seq reads remains a challenging task. Over recent years methods for both *de novo* and reference guided transcript reconstruction have accumulated rapidly. When combined with the large number of RNA-seq mapping tools deciding on the optimal transcriptome assembly strategy for a given organism and RNA-Seq data set (stranded/unstranded, polyA/ribodepleted) can be bewildering. In this article we showed that different assembly tools are complementary to each other; fully-reconstructing genes only partially reconstructed or missing entirely from alternative approaches. Similarly, when analysing multiple RNA-Seq samples, the complete transcript catalogue is often only obtained by collating together different assemblies. For a gene annotation project it is therefore typical to have multiple sets of transcripts, be they derived from alternative assemblers, different assembly parameters or arising from multiple samples. Our tool, Mikado, provides a framework for integrating transcript assemblies exploiting the inherent complementarity of the data to to produce a high-quality transcript catalogue.

Rather than attempting to capture all transcripts, our approach aims to mimic the selective process of manual curation by evaluating and identifying a subset of transcripts from each locus. The criteria for selection can be configured by the user, enabling them to for example to penalise gene models with truncated ORFs, those with non-canonical splicing, targets for nonsense mediated decay or chimeric transcripts spanning multiple genes. Such gene models may represent bona fide transcripts (with potentially functional roles), but can also arise from aberrant splicing or, as seen from our simulated data, from incorrect read alignment and assembly. Mikado acts as a filter principally to identify coding transcripts with complete ORFs and is therefore in line with most reference annotation projects that similarly do not attempt to represent all transcribed sequences. Our approach is made possible by integrating the data on transcript structures with additional information generally not utilised by transcript assemblers such as similarity to known proteins, the location of open reading frames and information on the reliability of splicing junctions. This information aids Mikado in performing operations such as discarding spurious alternative splicing events, or detecting chimeric transcripts. This allows Mikado to greatly improve in precision over the original assemblies, with in general minimal drops in recall. Moreover, similarly to TACO, Mikado is capable of identifying and resolving chimeric assemblies, which negatively affect the precision of many of the most sensitive tools, such as StringTie or the two meta-assemblers Cuffmerge and StringTie-merge.

Our experiments show that Mikado can aid genome annotation by generating a set of high quality transcript assemblies across a range of different scenarios. Rather than having to identify the best aligner/assembly combination for every project, Mikado can be used to integrate assemblies from multiple methods, with our approach reliably identifying the most accurate transcript reconstructions and allowing the user to tailor the gene set to their own requirements. It is also simple to incorporate assemblies from new tools even if the new method is not individually the most accurate approach. Given the challenges associated with short-read assembly it is desirable (when available) to integrate these with full-length cDNA sequences. Mikado is capable of correctly integrating analyses coming from different assemblers and technologies, including mixtures of Illumina and PacBio data. Our tool has already been employed for such a task on the large, repetitive genome of *Triticum aestivum* [31], where it was instrumental in selecting a set of gene models from over ten million transcript assemblies and PacBio IsoSeq reads. The consolidated dataset returned by Mikado was almost thirty times smaller than the original input dataset, and this polishing was essential both to ensure a high-quality annotation and to reduce the running times of downstream processes.

In conclusion, Mikado is a flexible tool which is capable of handling a plethora of data types and formats. Its novel selection algorithm was shown to perform well in model organisms and was central in the genome annotation pipeline of various species [31–33]. Its deployment should provide genome annotators with another powerful tool to improve the accuracy of data for subsequent *ab initio* training and evidence-guided gene prediction.

## Methods

### Input datasets

For *C. elegans, D. melanogaster* and *H. sapiens*, we retrieved from the European Nucleotide Archive (ENA) the raw reads used for the evaluation in [18], under the Bioproject PRJEB4208. We further selected and downloaded a publicly available strand-specific RNA-Seq dataset for *A. thaliana*, PRJEB7093. Congruently with the assessment in [18], we used genome assemblies and annotations from EnsEMBL v. 70 for all metazoan species, while for *A. thaliana* we used the TAIR10 version. For all species, we simulated reads using the input datasets as templates. Reads were trimmed with Trim-Galore v0.4.0 [34] and aligned onto the genome with Bowtie v1.1.2 [35] and HISAT v2.0.4 [7]. The HISAT alignments were used to calculate the expression levels for each transcript using Cufflinks v2.2.1, while the Bowtie mappings were used to generate an error model for the SPANKI Simulator v.0.5.0 [36]. The transcript coverages and the error model were then used to generate simulated reads, at a depth of 10X for *C. elegans* and *D. melanogaster* and 3X for *A. thaliana* and *H. sapiens*. A lower coverage multiplier was applied to the latter species to have a similar number of reads for all four datasets, given the higher sequencing depth in the *A. thaliana* dataset and the higher number of reference transcripts in H. sapiens. cDNA sequences for *A. thaliana* were retrieved from the NCBI Nucleotide database on the 21st of April 2017, using the query:

~~~
“Arabidopsis” [Organism] OR arabidopsis[All Fields]) AND “Arabidopsis thaliana”[porgn] AND biomol\_mrna [PROP]
~~~

For the second experiment on *H. sapiens*, we sequenced two samples of the Stratagene Universal Human Reference RNA (catalogue ID#740000), which consists of a mixture of RNA derived from ten different cell lines. One sample was sequenced on an Illumina HiSeq2000 and the second on a Pacific Biosciences RSII machine. Sequencing runs were deposited in ENA, under the project accession code PRJEB22606.

#### Preparation and sequencing of Illumina libraries

The libraries for this project were constructed using the NEXTflex Rapid Directional RNA-Seq Kit (PN: 5138-08) with the NEXTflex^™^ DNA Barcodes - 48 (PN: 514104) diluted to 6μM. The library preparation involved an initial QC of the RNA using Qubit DNA (Life technologies Q32854) and RNA (Life technologies Q32852) assays as well as a quality check using the PerkinElmer GX with the RNA assay (PN:CLS960010)

1 ¼g of RNA was purified to extract mRNA with a poly-A pull down using biotin beads, fragmented and first strand cDNA was synthesised. This process reverse transcribes the cleaved RNA fragments primed with random hexamers into first strand cDNA using reverse transcriptase and random primers. The second strand synthesis process removes the RNA template and synthesizes a replacement strand to generate dscDNA. The ends of the samples were repaired using the 3’ to 5’ exonuclease activity to remove the 3’ overhangs and the polymerase activity to fill in the 5’ overhangs creating blunt ends. A single ‘A’ nucleotide was added to the 3’ ends of the blunt fragments to prevent them from ligating to one another during the adapter ligation reaction. A corresponding single ‘T’ nucleotide on the 3’ end of the adapter provided a complementary overhang for ligating the adapter to the fragment. This strategy ensured a low rate of chimera formation. The ligation of a number indexing adapters to the ends of the DNA fragments prepared them for hybridisation onto a flow cell. The ligated products were subjected to a bead based size selection using Beckman Coulter XP beads (PN: A63880). As well as performing a size selection this process removed the majority of un-ligated adapters. Prior to hybridisation to the flow cell the samples were PCR’d to enrich for DNA fragments with adapter molecules on both ends and to amplify the amount of DNA in the library. Directionality is retained by adding dUTP during the second strand synthesis step and subsequent cleavage of the uridine containing strand using Uracil DNA Glycosylase. The strand that was sequenced is the cDNA strand. The insert size of the libraries was verified by running an aliquot of the DNA library on a PerkinElmer GX using the High Sensitivity DNA chip (PerkinElmer CLS760672) and the concentration was determined by using a High Sensitivity Qubit assay and q-PCR.

The constructed stranded RNA libraries were normalised and equimolar pooled into two pools. The pools were quantified using a KAPA Library Quant Kit Illumina/ABI (KAPA KK4835) and found to be 6.71 nM and 6.47 nM respectively. A 2nM dilution of each pool was prepared with NaOH at a final concentration of 0.1N and incubated for 5 minutes at room temperature to denature the libraries. 5 μl of each 2 nM dilution was combined with 995 μl HT1 (Illumina) to give a final concentration of 10 pM. 135 μl of the diluted and denatured library pool was then transferred into a 200 μl strip tube, spiked with 1 % PhiX Control v3 (Illumina FC-110-3001) and placed on ice before loading onto the Illumina cBot with a Rapid v2 Paired-end flow-cell and HiSeq Rapid Duo cBot Sample Loading Kit (Illumina CT-403-2001). The flow-cell was loaded on a HiSeq 2500 (Rapid mode) following the manufacturer’s instructions with a HiSeq Rapid SBS Kit v2 (500 cycles) (Illumina FC-402-4023) and HiSeq PE Rapid Cluster Kit v2 (Illumina PE-402-4002). The run set up was as follows: 251 cycles/7 cycles(index)/251 cycles utilizing HiSeq Control Software 2.2.58 and RTA 1.18.64. Reads in .bcl format were demultiplexed based on the 6bp Illumina index by CASAVA 1.8 (Illumina), allowing for a one base-pair mismatch per library, and converted to FASTQ format by bcl2fastq (Illumina).

#### Preparation and sequencing of PacBio libraries

The Iso-Seq libraries were created starting from 1 μg of human total RNA and full-length cDNA was then generated using the SMARTer PCR cDNA synthesis kit (Clontech, Takara Bio Inc., Shiga, Japan) following PacBio recommendations set out in the Iso-Seq method (http://goo.gl/1Vo3Sd). PCR optimisation was carried out on the full-length cDNA using the KAPA HiFi PCR kit (Kapa Biosystems, Boston USA) and 12 cycles were sufficient to generate the material required for ELF size selection. A timed setting was used to fractionate the cDNA into 12 individual sized fractions using the SageELF (Sage Science Inc., Beverly, USA), on a 0.75 % ELF Cassette. Prior to further PCR, the ELF fractions were equimolar pooled into the following sized bins: 0.7-2kb, 2-3kb, 3-5kb and ¿ 5kb. PCR was repeated on each sized bin to generate enough material for SMRTbell library preparation, this was completed following Pacbio recommendations in the Iso-Seq method. The four libraries generated were quality checked using Qubit Florometer 2.0 and sized using the Bioanalyzer HS DNA chip. The loading calculations for sequencing were completed using the PacBio Binding Calculator v2.3.1.1 (https://github.com/PacificBiosciences/BindingCalculator). The sequencing primer was used from the SMRTbell Template Prep Kit 1.0 and was annealed to the adapter sequence of the libraries. Each library was bound to the sequencing polymerase with the DNA/Polymerase Binding Kit v2 and the complex formed was then bound to Magbeads in preparation for sequencing using the MagBead Kit v1. Calculations for primer and polymerase binding ratios were kept at default values. The libraries were prepared for sequencing using the PacBio recommended instructions laid out in the binding calculator. The sequencing chemistry used to sequence all libraries was DNA Sequencing Reagent Kit 4.0 and the Instrument Control Software version was v2.3.0.0.140640. The libraries were loaded onto PacBio RS II SMRT Cells 8Pac v3; each library was sequenced on 3 SMRT Cells. All libraries were run without stage start and 240 minute movies per cell. Reads for the four libraries was extracted using SMRT Pipe v2.3.3, following the instructions provided by the manufacturer at https://github.com/PacificBiosciences/cDNA_primer.

### Alignments and assemblies

Reads from the experiments were aligned using STAR v2.4.1c and TopHat v2.0.14. For STAR, read alignment parameters for all species were as follows:

~~~
--outFilterMismatchNmax 4 --alignSJoverhangMin 12 --
    alignSJDBoverhangMin 12 --outFilterIntronMotifs
    RemoveNoncanonical --alignEndsType EndToEnd --
    alignTranscriptsPerReadNmax 100000 --
    alignIntronMin MININTRON --alignIntronMax
    MAXINTRON --alignMatesGapMax MAXINTRON
~~~

whereas for TopHat2 we used the following parameters:

~~~
-r 50 -p 4 --min-anchor-length 12 --max-multihits 20
    --library-type fr-unstranded -i MININTRON -I
    MAXINTRON
~~~

The parameters “MINTRON” and “MAXINTRON” were varied for each species, as follows:

- *A. thaliana*: minimum 20, maximum 10000
- *C. elegans*: minimum 30, maximum 15,000
- *D. melanogaster*: minimum 20, maximum 10,000
- *H. sapiens*: minimum 20, maximum 10,000

Each dataset was assembled using four different tools: CLASS v 2.12, Cufflinks 2.1.1, StringTie v. 1.03, and Trinity r20140717. Command lines for the tools were as follows:

- CLASS: we executed this tools through a wrapper included in Mikado, `class_xun.py`, with command line parameters `-F 0.05`
- Cufflinks: `-u -F 0.05`; for the *A. thaliana* dataset, we further specified `--library-type fr-firststrand`.
- StringTie: `-m 200 -f 0.05`
- Trinity: `--genome_guided_max_intron MAXINTRON` (see above)

Trinity assemblies were mapped against the genome using GMAP v20141229 [37], with parameters `-n 0 --min-trimmed-coverage=0.70 --min-identity=0.95` For simulated data, we elected to use a more modern version of Trinity (v.2.3.2) as the older version was unable to assemble transcripts correctly for some of the datasets. For assembling separately the samples in PRJBE7093, we used Cufflinks (v.2.2.1) and StringTie v1.2.3, with default parameters.

### Mikado analyses

All analyses were run with Mikado 1.0.1, using Daijin to drive the pipeline. For each species, we built a separate reference protein dataset, to be used for the BLAST comparison (see Table ST4). We used NCBI BLASTX v2.3.0 [38], with a maximum evalue of 10e-7 and a maximum number of targets of 10. Open reading frames were predicted using TransDecoder 3.0.0 [9]. Scoring parameters for each species can be found in Mikado v1.0.1, at https://github.com/lucventurini/mikado/tree/master/Mikado/configuration/scoring_files, with a name scheme of species_name_scoring.yaml (eg. “athaliana_scoring.yaml” for *A. thaliana*). The same scoring files were used for all runs, both with simulated and real data. Filtered junctions were calculated using Portcullis v1.0 beta5, using default parameters.

Mikado was instructed to look for models with - among other features - a good UTR/CDS proportion (adjusted per species), homology to known proteins, and a high proportion of validated splicing junctions. We further instructed Mikado to remove transcripts that do not meet minimum criteria such as having at least a validated splicing junction if any is present in the locus, and a minimum transcript length or CDS length. The configuration files are bundled with the Mikado software as part of the distribution.

### Details on the algorithms of Mikado

The Mikado pipeline is divided into three distinct phases.

#### Mikado prepare

Mikado prepare is responsible for bringing together multiple annotations into a single GTF file. This step of the pipeline is capable of handling both GTF and GFF3 files, making it adaptable to use data from most assemblers and cDNA aligners currently available. Mikado prepare will not just uniform the data format, but will also perform the following operations:

1. It will optionally discard any model below a user-specified size (default 200 base pairs).
2. It will analyse the introns present in each model, and verify their canonicity. If a model is found to contain introns from both strands, it will be discarded by default, although the user can decide to override this behaviour and keep such models in. Each multiexonic transcript will be tagged with this information, making it possible for Mikado to understand the number of canonical splicing events present in a transcript later on.
3. Mikado will also switch the strand of multiexonic transcripts if it finds that their introns are allocated to the wrong strand, and it will strip the strand information from any monoexonic transcript coming from non-strand specific assemblies
4. Finally, Mikado will sort the models, providing a coordinate-ordered GTF file as output, together with a FASTA file of all the cDNAs that have been retained.

*Mikado prepare* uses temporary SQLite databases to perform the sorting operation with a limited amount of memory. As such, it is capable of handling millions of transcripts from multiple assemblies with the memory found on a regular modern desktop PC (lower than 8GB of RAM).

#### Mikado serialise

*Mikado serialise* is the part of the pipeline whose role is to collect all additional data on the models, and store it into a standard database. Currently Mikado is capable of handling the following types of data:

1. FASTAs, ie the cDNA sequences produced by Mikado prepare, and the genome sequence.
2. Genomic BED files, containing the location of trusted introns. Usually these are either output directly from the aligners themselves (eg the “junctions.bed” file produced by TopHat) or derived from the alignment using a specialised program such as Portcullis.
3. Transcriptomic BED or GFF3 files, containing the location of the ORFs on the transcripts. These can be calculated with any program chosen by the user. We highly recommend using a program capable of indicating more than one ORF per transcript, if more than one is present, as Mikado relies on this information to detect and solve chimeric transcripts. Both TransDecoder and Prodigal have such capability.
4. Homology match files in XML format. These can be produced either by BLAST+ or by DIAMOND (v 0.8.7 and later) with the option “-outfmt 5”.

*Mikado serialise* will try to keep the memory consumption at a minimum, by limiting the amount of maximum objects present in memory (the threshold can be specified by the user, with the default being at 20,000). XML files can be analysed in parallel, so Mikado serialise can operate more efficiently if BLAST or DIAMOND runs are performed by pre-chunking the cDNA FASTA file and producing corresponding multiple output files.

Mikado serialise will output a database with the structure in SF8.

#### Mikado pick

Mikado pick selects the final transcript models and outputs them in GFF3 format. In contrast with many *ab initio* predictors, currently Mikado does not provide an automated system to learn the best parameters for a species. Rather, the choice of what types of models should be prioritised for inclusion in the final annotation is left to the experimenter, depending on her needs and goals. For the experiments detailed in this article, we configured Mikado to prioritise complete protein-coding models, and to apply only a limited upfront filtering to transcripts. A stricter upfront hard-filtering of transcripts, for example involving discarding any monoexonic transcript without sufficient homology support, might have yielded a more precise collated annotation at the price of discarding any potentially novel monoexonic genes. Although we provide the scoring files used for this paper in the software distribution, we encourage users to inspect them and adjust them to their specific needs. As part of the workflow, Mikado also produces tabular files with all the metrics calculated for each transcript, and the relative scores. It is therefore possible for the user to use this information to adjust the scoring model. The GFF3 files produced by Mikado comply with the formal specification of GFF3, as defined by the Sequence Ontology and verified using GenomeTools v.1.5.9 or later. Earlier versions of GenomeTools would not validate completely Mikado files due to a bug in their calculation of CDS phases for truncated models, see issue #793 on GenomeTools github: https://github.com/genometools/genometools/issues/793.

### Integration of multiple transcript assemblies

Evidential Gene v20160320 [23] was run with default parameters, in conjunction with CDHIT v4.6.4 [39]. Models selected by the tools were extracted from the combined GTFs using a mikado utility, mikado grep, and further clustered into gene loci using gffread from Cufflinks v2.2.1. StringTie-merge and Cuffmerge were run with default parameters. Limitedly to the experiment regarding the integration of assemblies from multiple samples, we used TACO v0.7. For all these three tools, we used their default isoform fraction parameter. The GTFs produced by the TACO meta-assemblies were reordered using a custom script (“sort_taco_assemblies.py”), present in the script repository.

### MAKER runs

We used MAKER v2.31.8 [40], in combination with Augustus 3.2.2 [41], for all our runs. GFFs and GTFs were converted to a match/match_part format for MAKER using the internal script of the tool “cufflinks2gff3.pl”. MAKER was run using MPI and default parameters; the only input files were the different assemblies produced by the tested tools.

### Comparison with reference annotations

All comparisons have been made using Mikado compare v1.0.1. Briefly, Mikado compare creates an interval tree structure of the reference annotation, which is used to find matches in the vicinity of any given prediction annotation. All possible matches are then evaluated in terms of nucleotide, junction and exonic recall and precision; the best one is reported as the best match for each prediction in a transcript map (TMAP) file. After exhausting all possible predictions, Mikado reports the best match for each reference transcript in the “reference map” (REFMAP) file, and general statistics about the run in a statistics file. Mikado compare is capable of detecting fusion genes in the prediction, defined as events where a prediction transcripts intersects at least one transcript per gene from at least two different genes, with either a junction in common with the transcript, or an overlap over 10% of the length of the shorter between the prediction or the reference transcripts. Fusion events are reported using a modified class code, with a “f,” prepending it. For a full introduction to the program, we direct the reader to the online documentation at https://mikado.readthedocs.io/en/latest/Usage/Compare.html.

#### Creation of reference and filtered datasets for the comparisons

For *A. thaliana*, we filtered the TAIR10 GFF3 to retain only protein coding genes. For the other three species, reference GTF files obtained through EnsEMBL were filtered using the “clean_reference.py” python script present in the “Assemblies” folder of the script repository (see the “Script availability” section). The YAML configuration files used for each species can be found in the “Biotypes” folder. The retained models constitute our reference transcriptome for comparisons.

For all our analyses, we deemed a transcript reconstructable if all of its splicing junctions (if any) and all its internal bases could be covered by at least one read. As read coverage typically decreases or disappears at the end of transcripts, we used the mikado utility “trim” to truncate the terminal UTR exons until their lengths reaches the maximum allowed value (50 bps for our analysis) or the beginning of the CDS section is found. BEDTools v. 2.27 beta (commit 6114307 [42]) was then used to calculate the coverage of each region. Detected junctions were calculated using Portcullis, specifically using the BED file provided at the end of Portcullis junction analysis step. The “get_filtered_reference.py” was then used to identify reconstructable transcripts.

For simulated datasets, we used the BAM file provided by SPANKI to derive the list of reconstructable transcripts. For the non-simulated datasets, we used the union of transcripts found to be reconstructable from each of the alignment methods. The utility “mikado util grep” was used to extract the relevant transcripts from the reference files. Details of the process can be found in the two snakemakes “compare.snakefile” and “compare_simulations.snakefile” present in the “Snakemake” directory of the script repository.

#### Calculation of comparison statistics

“Mikado compare” was used to assess the similarity of each transcript set against both the complete reference, and the reference filtered for reconstructable transcripts. Precision statistics were calculated from the former, while recall statistics were calculated from the latter.

### Script availability

Scripts and configuration files used for the analyses in this paper can be accessed at https://github.com/lucventurini/mikado-analysis.

### Customization and further development

Mikado allows to customize its run mode through the use of detailed configuration files. There are two basic configuration files: one is dedicated to the scoring system, while the latter contains run-specific details. The scoring file is divided in four different sections, and allows the user to specify which transcripts should be filtered out outright at any of the stages during picking, and how to prioritise transcripts through a scoring system. Details on the metrics, and on how to write a valid configuration file, can be found in the SI and at the online documentation (http://mikado.readthedocs.io/en/latest/Algorithms.html). These configuration files are intended to be used across runs, akin to how standard parameter sets are re-used in *ab initio* gene prediction programs, e.g. Augustus. The second configuration file contains parameters pertaining each run, such as the position of the input files, the type of database to be used, or the desired location for output files. As such, they are meant to be customised by the user for each experiment. Mikado provides a command, “mikado configure”, which will generate this configuration file automatically when given basic instructions.

## Declarations

### Ethics approval and consent to participate

Not applicable

### Consent for publication

Not applicable

### Availability of data and materials

The datasets supporting the conclusions of this article are included within the article (and its additional files). Transcript assemblies and gene annotation produced during the current study are available in a FigShare repository, together with the source code of the version of our software tool used to perform all experiments in this study, at this permanent address: https://figshare.com/projects/Leveraging_multiple_transcriptome_assembly_methods_for_improved_gene_structure_annotation/26149. Mikado is present on GitHub, at the address https://github.com/lucventurini/mikado. Many of the scripts used to control the pipeline executions, together with the scripts used to create the charts present in the article, can be found in the complementary repository https://github.com/lucventurini/mikado-analysis. All sequencing runs and reference sequence datasets used for this study are publicly available. Please see the section “Input datasets” for details.

### Authors’ contributions

LV and DS designed the Mikado algorithm and the pipeline. LV developed the code for the main program. LV and SC performed the complementary analyses. DLM developed the code for the Daijin pipeline. SC, GGK and DS tested the software and analyzed the resulting annotation to help refine the algorithm. LV and DS wrote the manuscript.

## Acknowledgements

This research was supported in part by the NBI Computing infrastructure for Science (CiS) group.

## Additional Files

Additional file 1 — Supplemental Information

Additional information for the main article, including supplemental figures and tables.

Additional file 2 — Reconstruction statistics for the input methods

This Excel file contains precision, recall and F1 statistics for the various methods tested.

